# Genetic diversity and distribution of indigenous soybean-nodulating bradyrhizobia in the Philippines

**DOI:** 10.1101/337394

**Authors:** Maria Luisa Tabing Mason, Baby Lyn Cortez Tabing, Akihiro Yamamoto, Yuichi Saeki

## Abstract

The diversity of indigenous bradyrhizobia from soils collected at 11 locations in the Philippines was investigated using PSB-SY2 local soybean cultivar as the host plant. Polymerase Chain Reaction-Restriction Fragment Length Polymorphism (PCR-RFLP) treatment for 16S rRNA, 16S-23S rRNA internal transcribed spacer (ITS) region and `rpo`B housekeeping gene was performed primarily to detect the genetic variation among the 424 isolates collected. Then, sequence analysis of 16S rRNA, ITS region and `rpo`B gene was performed for the representative isolates. Majority of the isolates were classified under `Bradyrhizobium elkanii, B. diazoefficiens, B. japonicum, Bradyrhizobium` sp., and few isolates were related to `B. yuanmingense`. Genetic variations observed through PCR-RFLP and sequence analyses of the ITS region and `rpo`B gene generally occurred in `B. elkanii`, suggesting an occurrence of gene transfer. Shannon’s diversity index showed varied results with a lowest score of 0.00 and highest at 0.98 indicating a very diverse population of bradyrhizobia across the country. Among all the factors considered in this work, soil management such as period of flooding and some soil properties provided major influence on the distribution and diversity of soybean bradyrhizobia in the country. Thus, it is proposed that the major micro-symbiont of soybean in the Philippines are `B. elkanii` for non-flooded soils, then `B. diazoefficiens` and `B. japonicum` for flooded soils.

**Importance:** Agriculture production in the Philippines has been and is currently heavily dependent on chemical inputs with mainly rice or corn mono-cropping that it rendered the soil acidic and unproductive. Legume research in the country are mainly focused on plant varietal improvements and very few are aimed at understanding the ecological niche of rhizobia present in the soil. Since soybean has mutual relationship with rhizobia, this legume is a good fallow crop or a rotation crop after rice and corn to help build up the nitrogen stock in the soil. The significance of this research is the better understanding of the ecological niche of indigenous soybean bradyrhizobia, particularly in a tropical archipelago like the Philippines. This work was conceptualized with the utmost goal to increase soybean yield by harnessing and evaluating the indigenous rhizobia in the soil to make production more sustainable and human-friendly.

## INTRODUCTION

The genus `Bradyrhizobium` is a major micro-symbiont of soybean (`Glycine max` [L.] Merrill) that could form nodule with legumes through symbiosis. The symbiotic relationship between rhizobia and legume is a highly specific, complicated and energy-exhaustive process (50) which may lead to improved crop yield if efficient and sustained. It is considered that `Bradyrhizobium` is the predominant genus of rhizobia in the tropics (9). Tropical regions have diverse environmental gradients that could influence the diversity of organisms. In this regard, high diversity of soybean-nodulating rhizobia exist in tropical regions (7) than temperate regions. By far, diverse species of bradyrhizobia were reported to be micro-symbionts of soybean such as `Bradyrhizobium japonicum, B. elkanii, B. yuanmingense, B. liaoningense, B. huanghuaihaiense` and `B. diazoefficiens` (3, 8, 16, 20, 34, 51, 55). Accordingly, many reports stated that soil pH, salinity, climate and cultural management influence the diversity and distribution of soybean bradyrhizobia (1, 5, 11, 22, 24, 37, 44, 54). High diversity of microorganisms in the soil plays an important role in maintaining soil health and vast, accurate information about this could lead to better crop productivity.

Philippines is a tropical country characterized by high temperature, humidity and abundant rainfall. It has two pronounced seasons based mainly from rainfall: (1) dry season – from December to May and (2) rainy season – from June to November. Aside from Baguio City, the average temperature in the country is almost similar at 25.5 – 27.5°C. The country’s agricultural production system has been heavily dependent on chemical inputs with rice or corn mono-cropping that it rendered the soil acidic and unproductive. Thus, the need for an effective and efficient symbiosis between legume and rhizobia became more important to increase crop’s yield in a sustainable and environmentally safe production system. Soybean is a good fallow or rotation crop after rice and corn. Although rice is the major agricultural product in the Philippines, recent trend recognized the role of soybean in nutrition and soil fertility restoration. This prompted its Government to create a Research and Development Roadmap for Soybean to increase the area and volume of production. This also paved the way for researchers to improve inoculation techniques to support the government’s endeavor.

Yet, inoculation does not always succeed due to several reasons such as incompatibility between the macro and micro-symbiont (12, 52, 53), competition between the indigenous and inoculated rhizobia (15, 42) and the above stated factors that influence the ecology and diversity of rhizobia. Hence, it is necessary to understand the factors influencing the ecology and diversity of indigenous rhizobia prior to inoculation.

Except for one location in the Philippines, which is Nueva Ecija (24), there is no information about the indigenous soybean rhizobia in Philippine soils. Since the Philippines is composed of more than 7,000 islands surrounded by bodies of water with considerable variation in agro-environmental conditions, it is hypothesized that diverse species of bradyrhizobia can be collected and identified to help in improving future inoculation techniques. Therefore, soil samples were collected from 11 locations in the Philippines to obtain information about the diversity and distribution of indigenous soybean bradyrhizobia. A Philippine soybean cultivar was used to trap the indigenous rhizobia from the collected soils then, PCR-RFLP and sequence analysis of the targeted genes were performed.

## MATERIALS AND METHODS

### Soil Collection

Soil samples were collected from eleven locations previously/currently planted with soybean and/or other legumes (Fig. 1). Information on the study sites are summarized in Table 1. Surface litters were removed before obtaining a bar of soil with dimension of 20 cm depth and 2 to 3 cm thickness and weighed approximately 1 kg. The 1 kg of composite soil sample per location was obtained from mixing and quartering at least 5 sub-samples that weighed approximately 1kg each. Half of the 1 kg composite soil was air dried and pulverized for soil analyses that include pH and electrical conductivity (EC) by 1:2.5 water extraction method, Total C, Total N, Bray P, K (flame spectrophotometer) and soil texture then, the remaining 0.5 kg was freshly used for soybean cultivation. Soil texture was analyzed by Hydrometer method as described previously (4), whereas C and N were analyzed by automatic high sensitive NC Analyzer Sumigraph NC-220F (Sumika Chemical Analysis Service. Ltd., Tokyo, Japan). Data of annual average temperature and rainfall were obtained from Philippine Atmospheric Geophysical and Astronomical Services Administration (PAGASA) website (https://www1.pagasa.dost.gov.ph/) which were averages from the last two decades.

**Table 1.** Agro-environmental information on the location of the study sites. All chemical and physical properties of the soil are analyzed in this study.

| Location | Coordinate | pH | EC<br>(dS/m) | C<br>(%) | N<br>(%) | Bray<br>P<br>(ppm) | K<br>(ppm) | Sand<br>(%) | Silt<br>(%) | Clay<br>(%) | Ave. Temp.<br>(°C) | Flooding<br>Period<br>(month) |
| --- | --- | --- | --- | --- | --- | --- | --- | --- | --- | --- | --- | --- |
| Ilagan, Isabela **(IS)** | 17.30°N,122.01°E | 5.90 | 0.08 | 1.34 | 0.13 | 1.86 | 51.80 | 28.0 | 34.5 | 37.5 | 26.9 | 5 |
| Gamu, Isabela **(GI)** | 17.08°N,121.79°E | 5.52 | 0.15 | 1.85 | 0.17 | 2.30 | 58.60 | 28.2 | 33.8 | 38.0 | 27.0 | 10 |
| Baguio, Benguet **(BA)** | 16.40°N,120.60°E | 5.22 | 0.20 | 3.10 | 0.24 | 22.22 | 51.00 | 19.0 | 41.4 | 39.6 | 19.3 | 10 |
| Nueva Ecija1 **(NE1)** | 15.74°N,120.93°E | 6.21 | 0.05 | 1.37 | 0.13 | 6.74 | 73.90 | 28.7 | 34.6 | 36.7 | 26.9 | 5 |
| Nueva Ecija2 **(NE2)** | 15.74°N,120.93°E | 5.81 | 0.12 | 2.36 | 0.22 | 21.63 | 49.40 | 27.4 | 34.7 | 37.9 | 26.9 | 10 |
| Irosin, Sorsogon **(SO)** | 12.72°N,124.04°E | 5.26 | 0.15 | 1.92 | 0.22 | 2.57 | 55.80 | 28.9 | 33.6 | 37.5 | 27.1 | 10 |
| Abuyog, Leyte **(LT)** | 10.67°N,125.04°E | 5.80 | 0.12 | 1.50 | 0.15 | 6.39 | 174.20 | 29.2 | 32.8 | 38.0 | 27.0 | 10 |
| La Carlota, Negros Occidental **(NR)** | 10.24°N,122.59°E | 5.62 | 0.15 | 0.63 | 0.07 | 20.44 | 74.10 | 28.0 | 34.2 | 37.8 | 27.7 | 5 |
| Ubay, Bohol **(BO)** | 9.99°N,124.45°E | 5.82 | 0.11 | 0.63 | 0.06 | 2.80 | 47.80 | 29.7 | 33.5 | 36.8 | 27.2 | 5 |
| Sultan Kudarat, Maguindanao **(SK)** | 6.51°N,124.42°E | 6.64 | 0.14 | 2.48 | 0.19 | 4.53 | 59.60 | 24.0 | 34.1 | 41.9 | 27.3 | 10 |
| Tupi, South Cotabato **(SC)** | 6.34°N,124.97°E | 5.52 | 0.15 | 1.36 | 0.14 | 31.18 | 47.20 | 19.5 | 42.5 | 38.0 | 25.4 | 10 |

**Figure 1.**
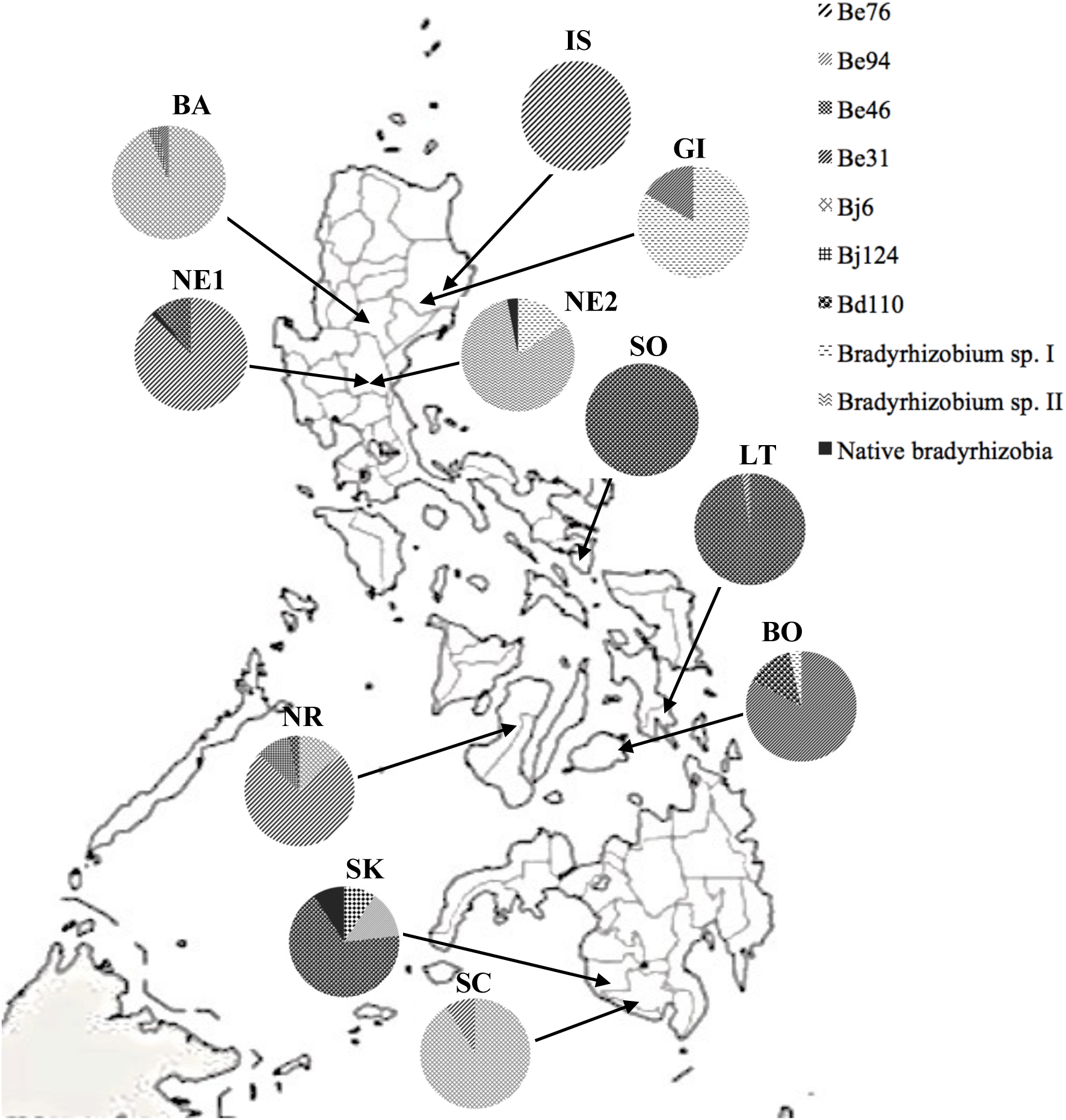
Distribution of soybean-nodulating bradyrhizobia in the Philippines from the result of Restriction Fragment Length Polymorphism (RFLP) and sequence analysis of the 16S-23S rRNA gene ITS region. (map from http://www.freeusandworldmaps.com).

### Isolation of indigenous soybean rhizobia

A local cultivar PSB-SY2, which is a commonly available soybean cultivar in the Philippines, was used to isolate the indigenous soybean rhizobia. Surface-sterilized soybean seeds were planted in 1-liter culture pots (n = 4). Sterilization of soybean seeds was done by soaking in 70% ethanol and sodium hypochlorite solution as previously described (34) prior to planting. Culture pots were filled with vermiculite containing N-free nutrient solution (33) at 40% (vol/vol) distilled water content and were autoclaved for 20 min at 121°C. Soil sample (2 to 3 g) was placed on the vermiculite at a depth of 2 to 3 cm, the seeds were then sown on the soil, and the pot was weighed. Plants were grown for 4 weeks in growth chamber (33°C for 16 h, day; 28°C for 8h, night), and were supplied weekly with sterile distilled water until the initial weight of the pot was reached.

After 4 weeks, 15 to 20 nodules were randomly collected from the roots of each soybean plant and sterilized with ethanol and sodium hypochlorite solution as described previously (47). Each nodule was homogenized in sterile distilled water, streaked onto a yeast extract mannitol agar (YMA; 49) plate medium, and incubated for about 1 week in the dark at 28°C. A single colony was streaked onto YMA plate containing 0.002% (wt/wt) bromothymol blue (BTB; 17) to determine the genus then, incubated as described above. Pure single colonies were obtained by repeated streaking into YMA plates.

All the isolates obtained from the 11 locations were used for 16S rRNA RFLP analysis to determine the genus of representative isolates for each operational taxonomic units (OTU). Based from the OTUs obtained, inoculation test was performed to determine their nodulation capability. Each isolate was cultured in YM broth culture (49) for about 1 week at 28°C, and the cultures were diluted with sterile distilled water to approximately 10^6^ cells ml^-1^. The soybean seeds were sown as described above but without soil, and inoculated with a 1-ml aliquot of each isolate per seed, replicated thrice. Nodule formation was assessed after 4 weeks in a growth chamber under similar conditions described above. A control pot (un-inoculated) was also prepared under similar conditions.

### DNA extraction

The pure single colony isolated from the YMA plate from repeated streaking was cultured in HEPES-MES (HM) broth culture (6, 40) for 3 to 4 days at 28°C with continuous agitation at 120 rpm. Thereafter, the bacteria cells cultured in the HM broth were collected by centrifugation and washed with sterile distilled water. Extraction of DNA was done by using BL buffer as described (25) from the method reported by Hiraishi et al. (13).

### PCR-RFLP of 16S rRNA gene, ITS region, and rpoB gene

Amplification of target genes was conducted using `Ex Taq` DNA polymerase (TaKaRa Bio, Otsu, Shiga, Japan) with primers and PCR cycle conditions previously used (24). The RFLP analyses of the 16S rRNA and ITS region were performed using the restriction enzymes `Hae`III, `Hha`I, `Msp`I and `Xsp`I (TaKaRa Bio) whereas for `rpo`B gene, enzymes `Hae`III, `Msp`I and `Alu`I (TaKaRa Bio) were used. `Bradyrhizobium` strains `B. japonicum` USDA 4, 6^T^, 38, 122, 123, 124, 129, 135, `B. diazoefficiens` USDA 110 ^T^`B. elkanii` USDA 31, 46, 76^T^, 94, and 130 and `B. liaoningense` USDA 3622^T^ (33) were used as reference strains. A 5.0 µl aliquot of the PCR product was digested with the restriction enzymes at 37°C for 16 h in a 20 µl reaction mixture. The restriction fragments were separated on 3 or 4% agarose gels in TBE buffer by means of electrophoresis and visualized with ethidium bromide.

### Sequence analysis

Based from the dendrogram of 16S rRNA, ITS region and `rpo`B gene constructed with Ward.D2 method of the R software v. 3.4.0, 31 representative isolates were selected for sequence analysis. The PCR amplified products of 31 representative isolates for ITS region and `rpo`B gene were purified according to the protocol of NucleoSpin^®^ Gel and PCR Clean-up (Macherey-Nagel, Germany). The DNA concentration of the purified product was determined by using NanoDrop 2000 Spectrophotometer (Thermo Scientific, U.S.A.). Preparation of samples from purified DNA followed the protocol for the premixed template and primer of the manufacturer (EUROFINS GENOMICS) using the previously designed sequence primers (24). After preparation, premixed samples were sent to the company for sequence analysis. After the ITS region-`rpo`B gene type of the 31 isolates were determined, 19 samples were randomly selected to confirm the sequences of 16S rRNA gene.

### Sequence alignment and construction of phylogenetic trees

Basic Local Alignment Search Tool (BLAST) program in DNA Databank of Japan (DDBJ) was used to determine the nucleotide homology. Sequences of type strains having similarity with our isolates of at least 99% for 16S rRNA, 96% for ITS region and 98% for `rpo`B gene were retrieved from BLAST database. The phylogenetic trees also included previously determined sequences of 16S rRNA and ITS region `Bradyrhizobium` genospecies (32, 48). Alignment of sequences obtained were performed using ClustalW. Phylogeny was determined by the Neighbor-Joining (38) method for the 16S rRNA, ITS region and `rpo`B gene. Genetic distances were calculated using Kimura 2-parameter model (18) in the Molecular Evolutionary Genetic Analysis (MEGA v7) software (19). Phylogenetic trees were bootstrapped with 1,000 replications of each sequence to evaluate the reliability of the tree topology. All the nucleotide sequences determined in this study were deposited in DDBJ at http://www.ddbj.nig.ac.jp/.

### Cluster and diversity analysis of indigenous bradyrhizobia

Only those isolates with reproducible fragments longer than 50bp in the electrophoresis gels were used for cluster analysis. The genetic distance between pairs of isolates (`D`) was calculated using the equation `D`_AB_ = 1 [2`N`_AB_/ (`N`_A_ + `N`_B_)], where `N`_AB_ represents the number of RFLP bands shared by strains A and B whereas `N`_A_ and `N`_B_ represent the numbers of RFLP bands found only in strains A and B, respectively (27, 39). The dendrogram were constructed by Ward.D2 method in R software v.3.4.0. Diversity analysis was performed by Shannon-Wiener diversity index as described previously (23, 29, 35). To expound on the community structure of dominant soybean bradyrhizobia, multi-dimensional scaling (MDS) analysis using Bray-Curtis Index was employed also in R software.

### Principal component analysis (PCA)

To detect the relationship between the agro-environmental factors and the distribution of soybean bradyrhizobia, PCA was performed in R software. The variables for the principal component include soil chemical properties, soil texture and some environmental data.

## RESULTS

### Isolation of indigenous bradyrhizobia

A total of 771 isolates was obtained from the study sites with a range of 63 to 79 isolates per location and were all used for the primary16S rRNA gene RFLP analysis. Samples were labeled with the combination of the abbreviation of the sampling site (IS - Ilagan; GI - Gamu; BA - Baguio; NE1 - 1^st^ location in Nueva Ecija; NE2 - 2^nd^ location in Nueva Ecija; SO - Sorsogon; LT - Leyte; BO - Bohol; NR - Negros; SK - Sultan Kudarat; and SC - South Cotabato) and the number of the isolate (1-63 or 1-79) (e.g., for South Cotabato, SC – 1-74).

We obtained 502 (65.11%) slow-growing (7-8 days) and 269 (34.89%) fast-growing (2-3 days) isolates. All the slow-growers which produced alkaline or neutral reaction in YMA plates with BTB were considered as `Bradyrhizobium` (16). All the slow-growers produced nodules on soybean and were used for the ITS region amplification. We were able to obtain 424 `Bradyrhizobium` isolates that were successfully amplified at approximately 900bp amplicons using primers specific for bradyrhizobia.

### PCR-RFLP analysis of 16S rRNA, ITS region and rpoB gene

The 424 isolates were all used for the RFLP treatment of ITS region and `rpo`B gene. The phylogenetic trees according to the differences in fragment size and pattern from the RFLP treatment of the ITS region and `rpo`B gene are presented in Figures 2 and 3, respectively. In ITS region, eleven clusters were identified of which three clusters belong to `B. elkanii` (Be46, Be76, Be94), one cluster to `B. diazoefficiens` (Bd110), two clusters to `B. japonicum` (Bj6, Bj124), and five clusters were assigned to `Bradyrhizobium` species. The five clusters of `Bradyrhizobium` sp. consisted of dominant isolates from GI (30 out of 36) and NE2 (31 out of 35), and very minor isolates from NE1 (6 out of 56) and SK (4 out of 43). Meanwhile, ten clusters were classified for `rpo`B gene of which three are assigned to `B. elkanii` (Be46, Be94, Be130), one for `B. diazoefficiens` (Bd110), two for `B. japonicum` (Bj6, Bj124) and four to `Bradyrhizobium` species. It is noticeable that the isolate NR-60 which was clustered under Be76 in the ITS region became clustered under `Bradyrhizobium` sp. in the `rpo`B gene. The phylogenies of ITS region and `rpo`B gene showed genetic variations mostly for isolates grouped under `B. elkanii.`

**Figure 2.**
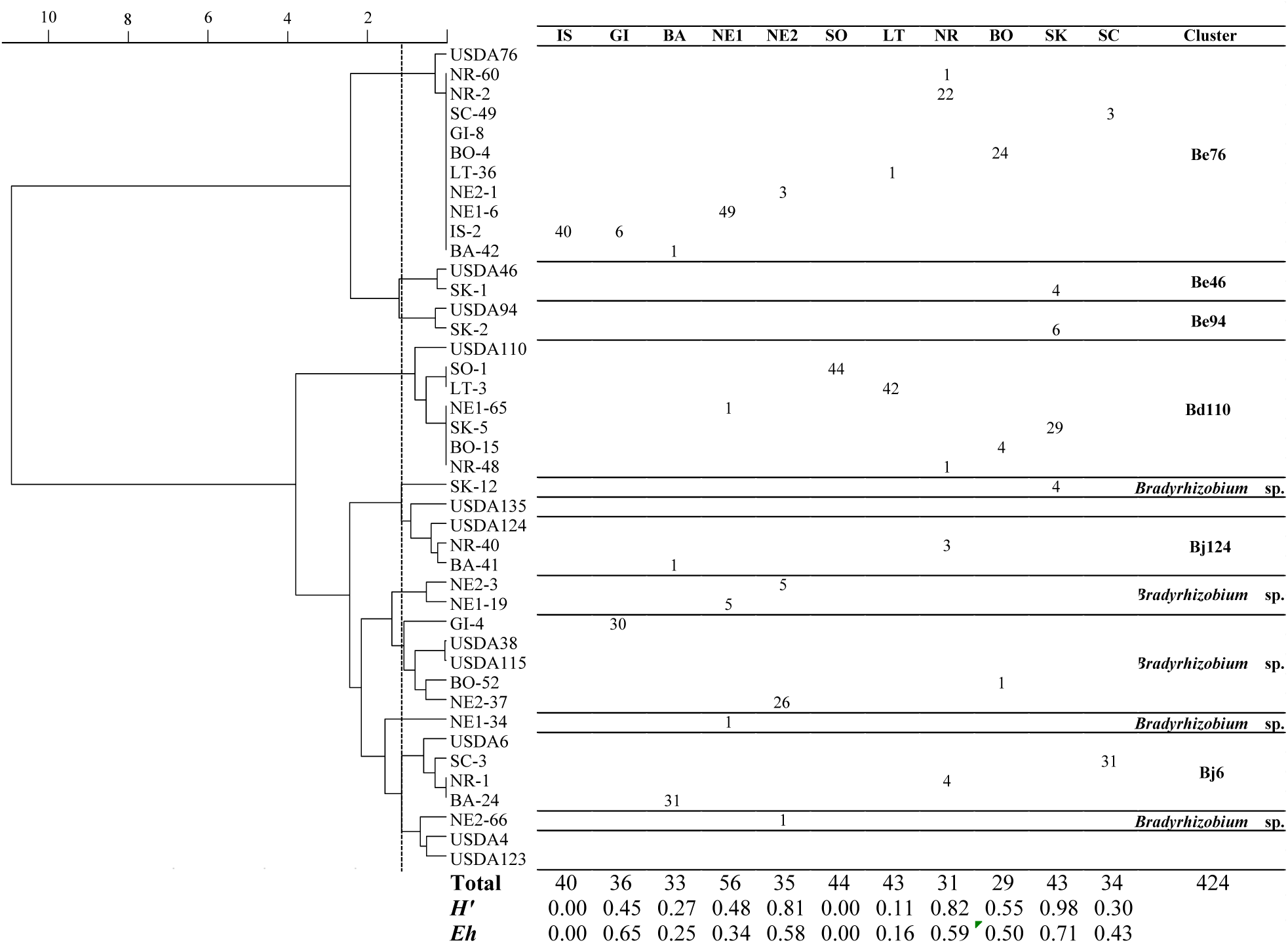
Dendrogram of the 16S-23S rRNA gene ITS region of soybean-nodulating bradyrhizobia in the Philippines with selected `Bradyrhizobium` USDA reference strains indicating the cluster distribution. The phylogenetic tree was constructed from the Ward.D2 method in R software v. 3.4.0. A similarity from the height of 1.2 between isolate SK-12 and `B. japonicum` USDA124 was used to distinguish the clusters. Shannon’s diversity (`H’`) and equitability (`Eh`) indices were computed with the formulae (`H’`= -Σ`Pi` ln `Pi`; `Eh`= `H’`/ ln `S`).

**Figure 3.**
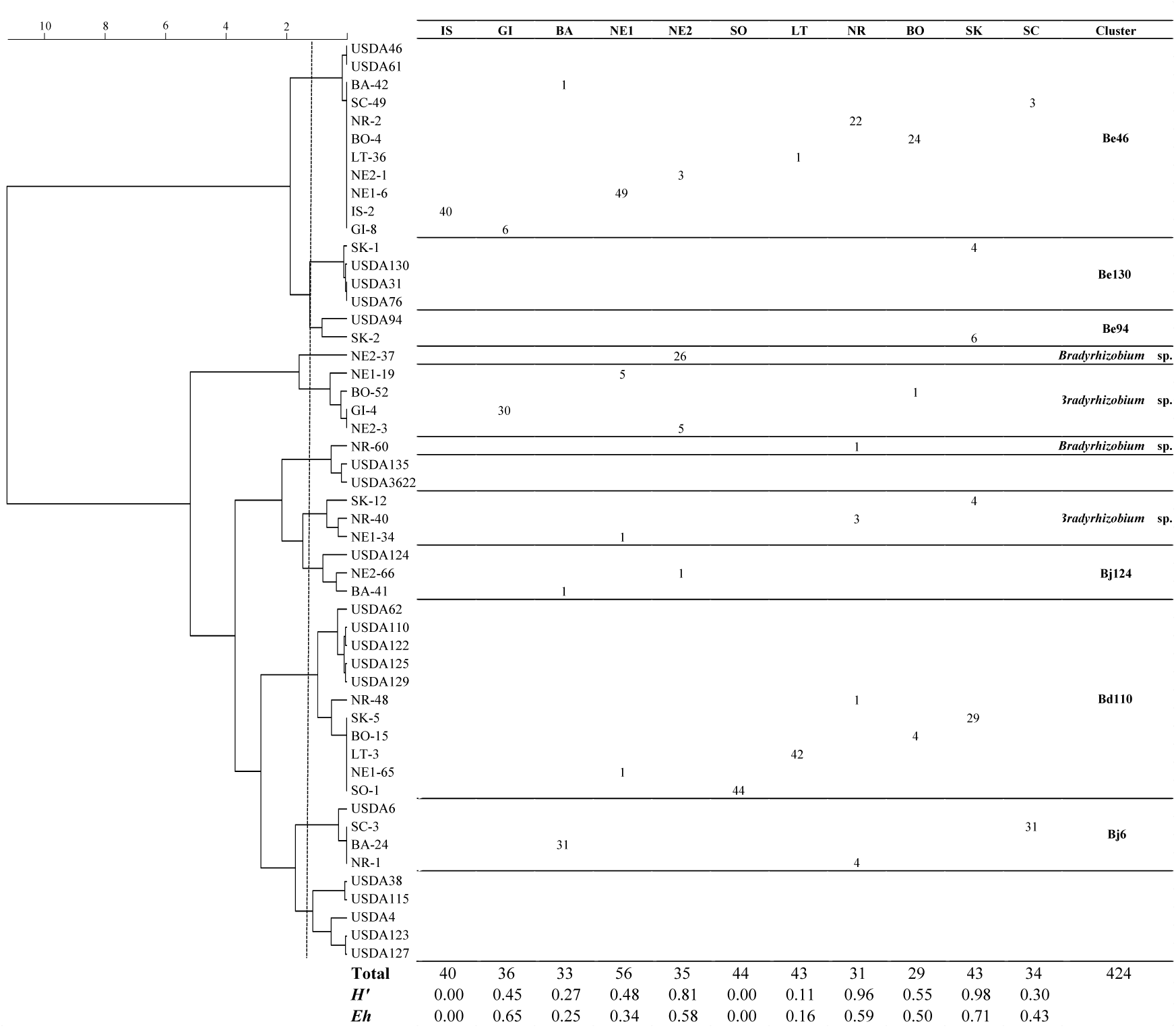
Dendrogram of the `rpo`B housekeeping gene of soybean-nodulating bradyrhizobia in the Philippines with selected `Bradyrhizobium` USDA reference strains indicating the cluster distribution. The phylogenetic tree was constructed from the Ward.D2 method in R software v.3.4.0. A similarity from the height of 1.2 between `B. elkanii` USDA130 and USDA94 was used to to distinguish the clusters. Shannon’s diversity (`H’`) and equitability (`Eh`) indices were computed with the formulae (`H’`= -Σ`Pi` ln `Pi`; `Eh`= `H’`/ ln `S`).

### Sequence analysis of 16S rRNA, ITS region and rpoB gene

Presented in Figure 4 are the phylogenetic trees from the result of sequence analysis. For ITS region (Fig. 4A), the isolates were predominantly grouped under four bradyrhizobia species which are `B. elkanii` (37.74%), `B. diazoefficiens` (28.54%), `B. japonicum` (16.51%), and `B. yuanmingense` (0.94%). A `Bradyrhizobium` sp. clade that makes up about 16.74% of the population was also observed including two independent single isolates NE1-34 and NE2-66. On the other hand, almost similar clusters were observed from `rpo`B gene (Fig. 4B) wherein eight delineations were distinguished with independent isolates NE1-34 and NR-40 that also belong to `Bradyrhizobium` sp.. Notably, genetic variations were detected on specific isolates between the ITS region and `rpo`B gene. For example, SK-1 belonged to Be46 cluster in ITS region but it was reclassified into Be130 cluster in `rpo`B gene. Moreover, the isolate NR-60 which belonged to Be cluster in the ITS region was noticeably under `Bradyrhizobium` sp. in `rpo`B gene. For all isolates belonging to Be clusters, only those that belong to Be94 remain unchanged from ITS region to `rpo`B gene (SK-2). No remarkable genetic diversity was observed for other clusters particularly for those under `B. diazoefficiens` and `B. yuanmingense`.

**Figure 4.**
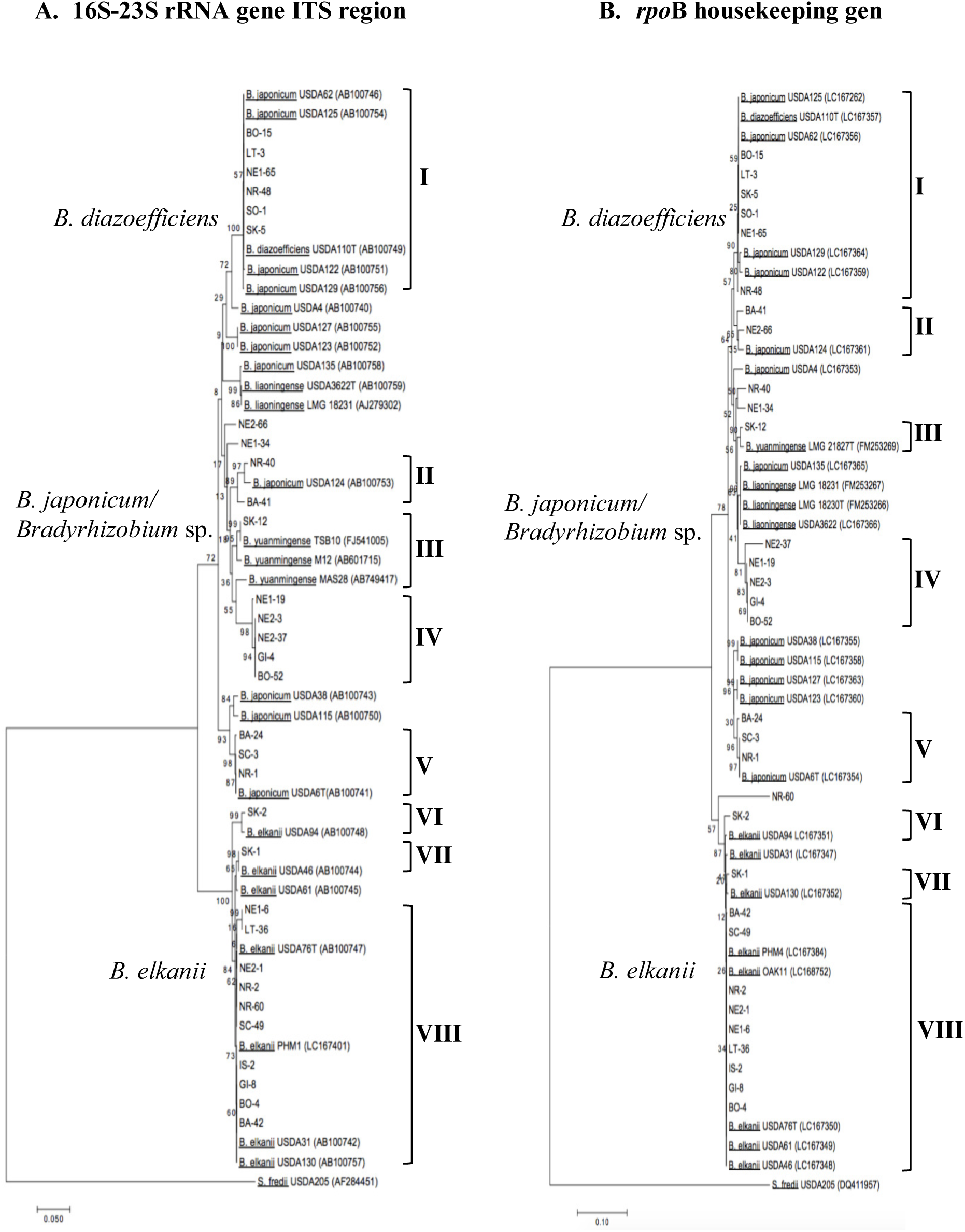
Phylogenetic tree based on sequence analysis of (A) 16S-23S rRNA gene ITS region and (B) `rpo`B housekeeping gene. The tree was constructed using the Neighbor-Joining method with the Kimura 2-parameter (K2P) distance correlation model and 1000 bootstrap replications in MEGA v.7 software. The accession numbers are indicated only for sequences obtained from BLAST. The isolates in this study are indicated with letters and number combinations, for example: **IS-2** – isolate no. 2 collected from Ilagan, Isabela.

### Cluster and diversity analysis

The clusters and diversity indices of soybean bradyrhizobia in the Philippines are presented in Figure 2. Isolates clustered to `B. elkanii` were present in almost all the locations except in Sorsogon (SO), wherein all the isolates were classified under `B. diazoefficiens`. Diversity and equitability indices obtained from the analysis of the ITS region (Fig. 2) were highest at Sultan Kudarat (SK) (`H’`=0.98, `Eh`=0.71), followed by Negros Occidental (NR) (`H’`=0.82, `Eh`=0.59). The lowest indices (`H’`=0.00, `Eh`=0.00) were obtained at Ilagan, Isabela and Sorsogon where all isolates belonged to `B. elkanii` and `B. diazoefficiens`, respectively. For `rpo`B gene (Fig. 3), diversity indices were still highest at the same location but a slight change was noticed in Negros Occidental (NR) which showed higher indices than in ITS region due to genetic variation detected from the isolate NR-60.

Meanwhile, the MDS plots clearly indicate the community structure and population dominance of each `Bradyrhizobium` species in accordance with their clusters and respective location (Fig. 5). The different clusters of `Bradyrhizobium` sp. obtained from in the phylogenetic trees of the ITS region and `rpo`B gene are clustered together in the plots (GI, NE2). Conversely, SO, LT and SK were grouped together because these locations are dominated by those clustered to `B. diazoefficiens`. Similar observations were obtained for the locations dominated by `B. elkanii` (IS, BO, NR, NE1) and `B. japonicum` USDA6 (BA, SC).

**Figure 5.**
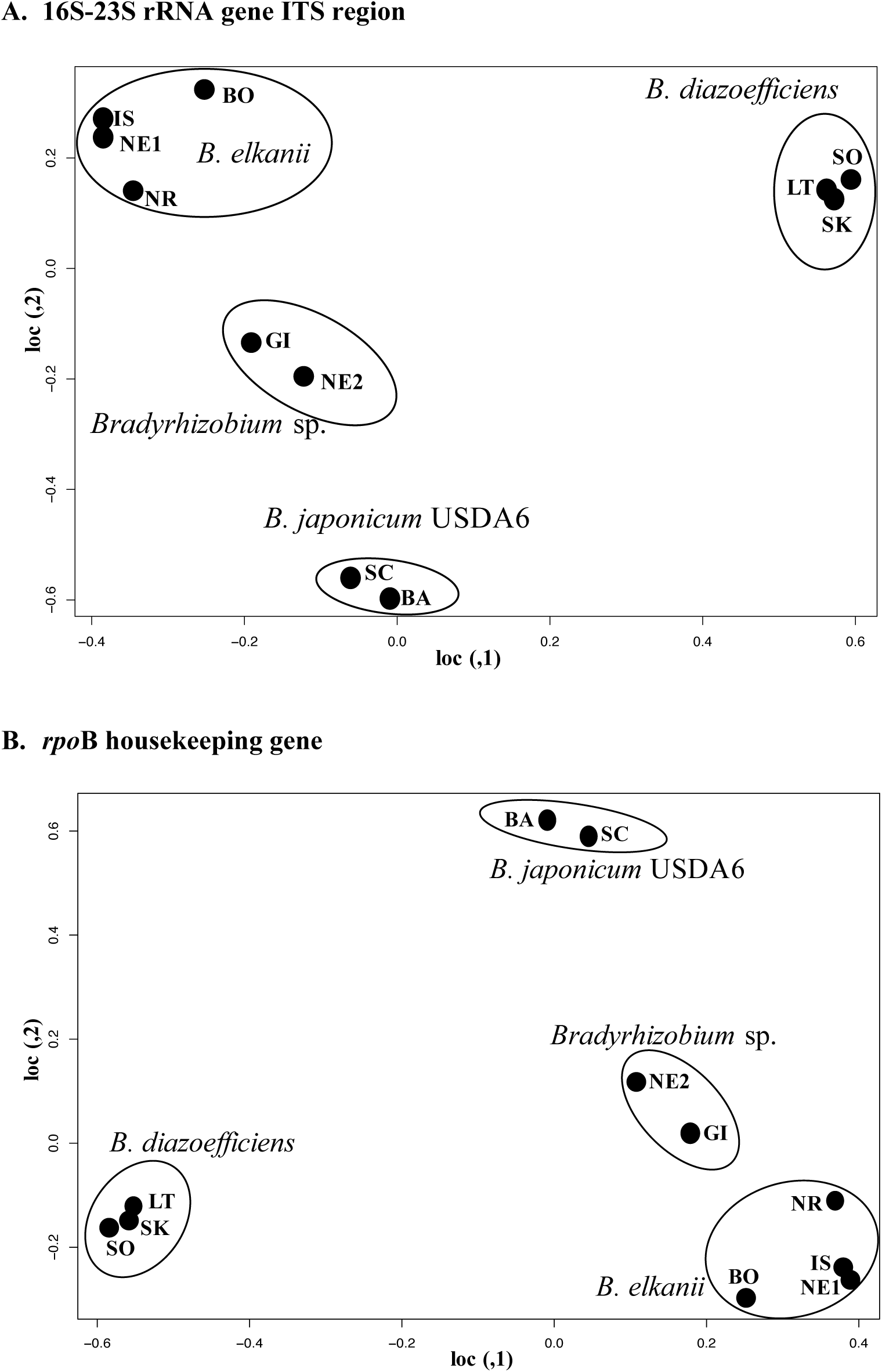
Community structure of indigenous soybean-nodulating bradyrhizobia in the Philippines showing the dominance of each `Bradyrhizobium` species at respective location. The figure was constructed using Bray-Curtis index through R software v.3.4.0. **A** – community structure from the 16S-23S rRNA gene ITS region. **B** – community structure from the `rpo`B gene.

### PCA of the factors affecting the diversity and distribution of bradyrhizobia

Presented in Figure 6 is the PCA plot showing the factors that influence the diversity and distribution of indigenous bradyrhizobia in the country. PC1 shows 47.74% proportion that is accounted for most of the variance and indicating the correlation between the parameters considered and the distribution of bradyrhizobia in specific locations. Strong influence of flooding period (FP), N, C, pH and clay content can be attributed to the abundance of `Bradyrhizobium` sp. and `B. diazoefficiens` USDA110^T^ while the dominance of `B. japonicum` USDA6^T^ can be attributed to the influence of high silt and P content in the soil. On the other hand, the abundance and distribution of `B. elkanii` is mainly correlated with the high temperature in the country and possibly higher amount of sand.

**Figure 6.**
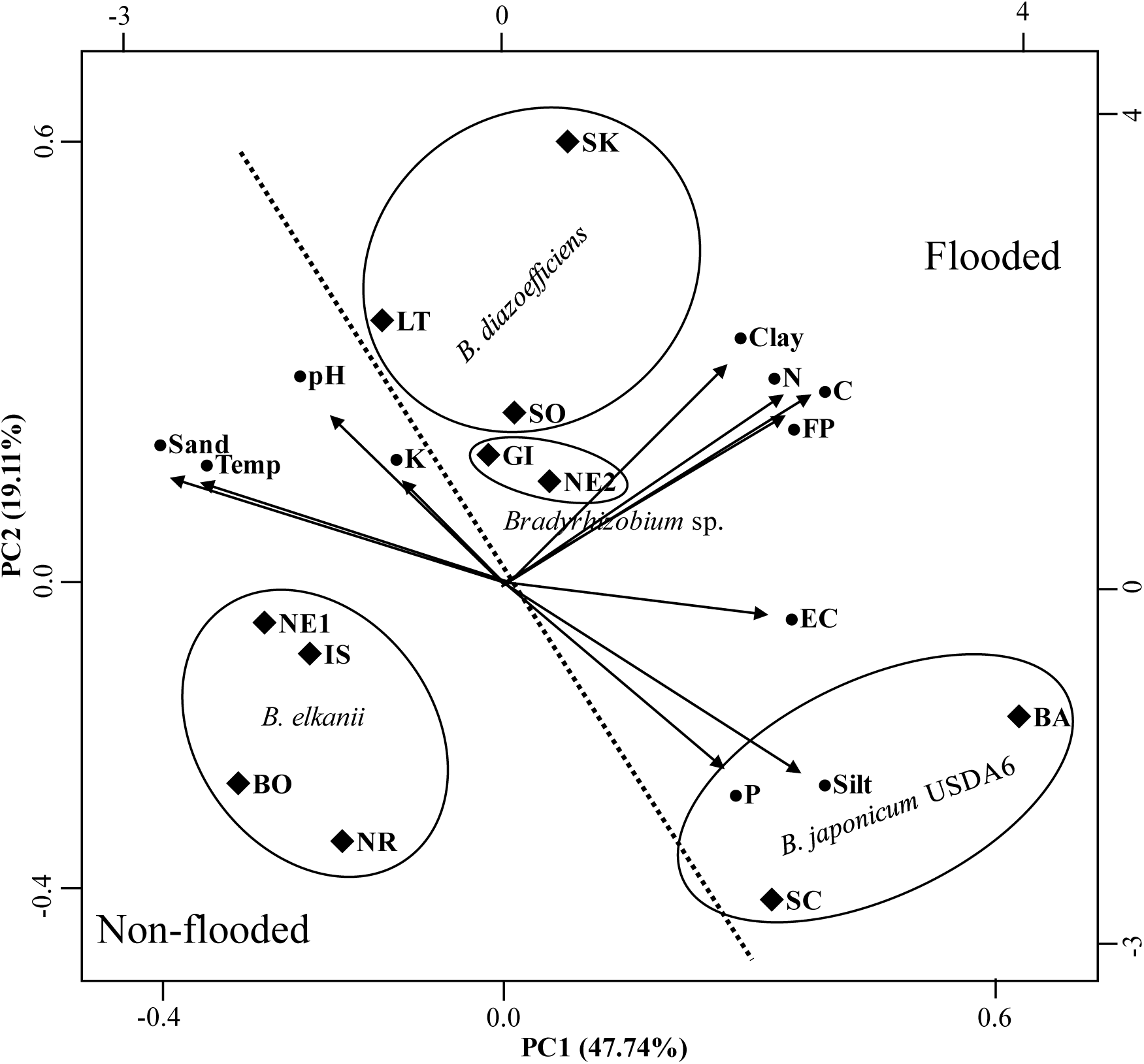
Principal Component Analysis (PCA) plot depicting the relationship between the dominance of `Bradyrhizobium` species in the respective locations and the agro-environmental factors considered in this study. • Blackened circle indicates the agro-environmental factors. ♦ Blackened diamond indicates the location of the soil sampling collection. FP-period of flooding. Dotted straight line indicates the separation between the flooded and non-flooded condition.

## DISCUSSION

### Geographic distribution and genetic diversity of bradyrhizobia

The high genetic diversity of indigenous soybean bradyrhizobia in the Philippines supported the hypothesis of this study. Aside from the high species diversity as revealed from the Shannon’s index, genetic diversity was also remarkable. Notably, most of the genetic variations were detected only on `B. elkanii` isolates suggesting that this species maybe easily influenced by recombination events and the occurrence of horizontal gene transfer as previously observed (24, 26). It was reported that genetic variations detected from different gene loci commonly occur in nature as a result of recombination events due to physical proximity, environmental changes and genetic distance (10). From the genome map of `B. elkanii` USDA76^T^ with a genome size of 9,484,767bp, `rpo`B gene (6,539,500bp) is located on almost opposite locus with the ITS region (411,600 – 410,900bp) as reported previously (30) so it may easily reflect genetic changes that occurred within the species.

In Nueva Ecija (NE), wherein the location of samples was almost similar in all aspect except for the soil water status, different species of bradyrhizobia dominated each location. NE1 isolates (dominated by `B. elkanii`) were collected from non-flooded condition while NE2 isolates (`Bradyrhizobium` sp.) were collected from flooded condition. Additionally, dominance of isolates clustered under `B. diazoefficiens` USDA110^T^ was observed in flooded soil and were generally distributed in the central islands (Sorsogon (SO), Leyte (LT), Negros Occidental (NR), Bohol (BO)) to southern region (Sultan Kudarat (SK)) of the Philippines. This signifies that cultural management such as waterlogging may affect the population dominance of strains belonging cluster Bd110. This result is supported by our previous reports where it was observed that the dominance of cluster `B. diazoefficiens` USDA110^T^ was enhanced by flooding condition in the soil (37) and that USDA110 cluster are dominant on fine-textured soil that are affected by water status and oxidation-reduction potential in the soil (36). Additionally, it was reported that the anaerobic condition in flooded soil of alluvial origin resulted in the dominance of `B. diazoefficiens` USDA110^T^ (43) which is also similar our results. On the other hand, a study stated that the diversity and abundance of `Bradyrhizobium` species were altered by cultural management and other soil-related properties (54). In our results, about 16% from the total population were classified under `Bradyrhizobium` sp. and these were found in both flooded and non-flooded soils. Interestingly, these strains are even dominant in two locations (GI and NE2) and mainly found in the Luzon island. We considered that these strains are novel species in the Philippines since the stated locations have no history of rhizobial inoculation and we did not obtain any highly similar sequences from the BLAST engine for its identification.

A minor population of strains related to `B. yuanmingense` (0.94%) was observed from South Cotabato (SK) and based from our result, the agro-environment factors considered have no strong influence on its identification. Several studies indicated that this bradyrhizobia is found in a variety of regions especially across Asia (3, 45), with increased population of `B. yuanmingense` in moderately acidic soils and cooler regions (1, 31). Our results revealed that isolates related to `B. yuanmingense` was found only on flooded soil with slight acidity.

### Factors affecting the diversity and distribution of bradyrhizobia

In this study, several agro-environmental factors were considered to investigate which parameter/s highly influence the distribution and genetic diversity of soybean bradyrhizobia in the Philippines. The high amount of silt and phosphorus in the soil positively influenced the distribution and dominance of Bj6 cluster in Baguio (BA) and South Cotabato (SK). The positive influence of phosphorus content on the abundance of `B. japonicum` sp., which were phylogenetically clustered with Bj6 was also observed earlier (54). Similarly, a previous report indicated that isolates clustered to `B. japonicum` USDA6^T^ are dominant in Andosols in Japan (43). Additionally, soils of South Cotabato (SC) are Andisols with mixed alluvium and sedimentary deposits where dominant strains of Bj6 cluster is found in this study. Since `B. japonicum` USDA6^T^ was reported to release N_2_O (41), which is one of the greenhouse gases, its impact in both agriculture and environment merits more attention for better understanding.

Meanwhile, the dominance of isolates clustered under `B. diazoefficiens` USDA110^T^ in Sorsogon (SO), Leyte (LT) and Sultan Kudarat (SK) can be explained by the periods of flooding in these areas as affected by high precipitation and soil management. These areas are usually planted with rice during wet season then, planted with rice and/or legume during dry season. In both season, rice farming is always done in waterlogged status. This provided an interesting insight particularly for Philippines agriculture since `B. diazoefficiens` USDA110^T^ was proven to be a highly effective and efficient inoculant for soybean (21, 46) including its complete denitrification ability (2, 14, 41, 43) that is beneficial for climate change mitigation. The locations where these strains were isolated had no history of USDA110 inoculation so these strains are considered indigenous. Further studies on the strains’ characteristics, possession of denitrification genes, and symbiotic relationship with different soybean cultivar in the Philippines will be helpful to test its potential as an efficient and effective inoculant.

As expected, the abundance and widespread distribution of `B. elkanii` in the Philippines is consistent with previous findings wherein this species is distributed in areas with slight to moderate acidity and sub-tropical to tropical region (1, 24, 34, 44). The species and genetic diversity of `B. elkanii` is the highest among all the other species in this report. However, it is worthy to note that aside from soil acidity and temperature, the non-flooded status of the soil as influenced by cultural practices indicates another major impact on the distribution of `B. elkanii`. This result might be one of the reasons for low yield of soybean across the country. It is known that `B. elkanii` provides lower N fixation and symbiotic efficiency (31) in comparison with `B. diazoefficiens` USDA110^T^. Additionally, `B. elkanii` is an NO_2_^-^ producer, and it is known that the interaction of nitrites to some soil components are related to environmental concerns. Thus, species of bradyrhizobia are important to conduct research on for its role in agriculture and environment.

The presence of `B. yuanmingense` as minor micro-symbionts of soybean in the Philippines added new information to existing literature especially for tropical bradyrhizobia.

We were able to determine that the major micro-symbionts of soybean in the Philippines are `B. elkanii` for non-flooded soils with higher soil temperature, `B. diazoefficiens` USDA110^T^ for fine-textured flooded soils and `B. japonicum` USDA6^T^ for flooded soils high in phosphorus and silt content. The considerable number of isolates classified as `Bradyrhizobium` sp. should be worthy specimen for further research because they might indicate evolution or adaptation mechanism of bradyrhizobia as influenced by land use changes. This observation was reported beforehand stating that land use system affected the diversity of species under `Bradyrhizobium` sp. (28). High genetic diversity of soybean bradyrhizobia was observed with possible indigenous strains that can be used for future studies. It is proposed that the distribution and genetic diversity of bradyrhizobia in the Philippines was mainly influenced by soil management (periods of flooding) and soil properties (soil type and other edaphic factors). This work was able to provide information that the ecology of bradyrhizobia is indeed complicated, particularly in a tropical region; and it depends on several abiotic and biotic factors.

## ACKNOWLEDGEMENTS

We would like to thank the following for their help during soil sampling: Ms. Jenny Castaneto of Department of Agriculture – Bureau of Agricultural Research (DA-BAR) of the Philippines; DA Regional offices in Baguio City and Bohol, Philippines; and Mr. Leolito Siase of Bureau of Soils and Water Management (BSWM), Philippines. This work was funded by JSPS KAKENHI (Grant-in-Aid for Scientific Research (C) No. 18K05376) and the Ministry of Education, Culture, Sports, Science and Technology (MEXT) Japanese Government Scholarship Program.

